# Piperacillin/tazobactam resistant, cephalosporin susceptible *Escherichia coli* bloodstream infections are driven by multiple acquisition of resistance across diverse sequence types

**DOI:** 10.1101/2020.09.18.302992

**Authors:** Thomas Edwards, Eva Heinz, Jon van Aartsen, Alex Howard, Paul Roberts, Caroline Corless, Alice J. Fraser, Christopher T. Williams, Issra Bulgasim, Luis E. Cuevas, Christopher M. Parry, Adam P. Roberts, Emily R. Adams, Jenifer Mason, Alasdair T. M. Hubbard

**Affiliations:** Department of Tropical Disease Biology, Liverpool School of Tropical Medicine, Pembroke Place, Liverpool, L3 5QA; Cent re for Drug and Diagnostics, Liverpool School of Tropical Medicine, Pembroke Place, Liverpool, L3 5QA; Department of Vector Biology, Liverpool School of Tropical Medicine, Pembroke Place, Liverpool, L3 5QA; Department of Clinical Sciences, Liverpool School of Tropical Medicine, Pembroke Place, Liverpool, L3 5QA; Wellcome Sanger Institute, Wellcome Genome Campus, Hinxton, CBl0 1SA; Liverpool University Hospital Foundation Trust, Prescot street, Liverpool, L7 8XP; Faculty of Science and Engineering, University of Wolverhampton, Wulfruna Street, Wolverhampton WV1 1LY; Alder Hey Children’s NHS Foundation Trust, Eaton Road, Liverpool, L12 2AP; Department of Biosciences, School of Science and Technology, Nottingham Trent University, Nottingham, NG11 8NS

## Abstract

Resistance to piperacillin/tazobactam (TZP) in *Escherichia coli* has predominantly been associated with mechanisms that confer resistance to third generation cephalosporins. Recent reports have identified *E. coli* strains with phenotypic resistance to piperacillin/tazobactam but susceptibility to third generation cephalosporins (TZP-R/3GC-S). In this study we sought to determine the genetic diversity of this phenotype in *E. coli* (*n* = 58) isolated between 2014-2017 at a single tertiary hospital in Liverpool, UK, as well as the associated resistance mechanisms. We compare our findings to a UK-wide collection of invasive *E. coli* isolates *(n* = 1509) with publicly available phenotypic and genotypic data. These data sets included the TZP-R/3GC-S phenotype *(n* = 68), a piperacillin/tazobactam and third generation cephalosporin-susceptible (TZP-S/3GC-S, *n* = 1271) phenotypes. The TZP-R/3GC-S phenotype was displayed in a broad range of sequence types which was mirrored in the same phenotype from the UK-wide collection, and the overall diversity of invasive *E. coli* isolates. The TZP-R/3GC-S isolates contained a diverse range of plasmids, indicating multiple acquisition events of TZP resistance mechanisms rather than clonal expansion of a particular plasmid or sequence type. The putative resistance mechanisms were equally diverse, including hyperproduction of TEM-1, either via strong promoters or gene amplification, carriage of inhibitor resistant β-lactamases, and an S133G *b/a*_CTX-M-15_ mutation detected for the first time in clinical isolates. Several of these mechanisms were present at a lower abundance in the TZP-S/3GC-S isolates from the UK-wide collection, but without the associated phenotypic resistance to TZP. Our findings highlight the complexity of this cryptic phenotype and the need for continued phenotypic monitoring, as well as further investigation to improve detection and prediction of the TZP-R/3GC-S phenotype from genomic data.

## Introduction

*Escherichia coli* is the most common cause of bacterial blood stream infections globally (1), accounting for 27% of all bacteraemic episodes, with a case fatality rate of 12% (2), and causing 78.8 blood stream infections per 100,000 people in the UK in 2014 (3). Antimicrobial resistance (AMR) in *E. coli* is increasingly prevalent (4–6) and extended spectrum β-lactamase (ESBL) production, mediating resistance to third generation cephalosporins (3GCs) and other β-lactam antibiotics (7), is of particular concern. ESBLs were recorded in approximately 11% of *E. coli* isolated from blood stream infections in the UK in 2018 (8).

One strategy to provide therapeutic options for antimicrobial resistant infections has been the combined use of β-lactamase inhibitors with β-lactam antibiotics to block the activity of β-lactamase enzymes, rendering the bacteria *de facto* susceptible (9). The inhibitor tazobactam, which inhibits class A β-lactamases and includes most ESBL enzymes, is commonly utilised in combination with the penicillin class antibiotic piperacillin (10). Tazobactam is a “suicide inhibitor”, as it irreversibly binds to β-lactamases, inactivating the enzyme (11). Piperacillin/tazobactam (TZP) has broad spectrum activity against Gram-negative and -positive bacteria (12), is well tolerated (13), available for paediatric use, and utilised in the UK as a first line empirical agent for serious infections, including pneumonia and intra-abdominal infections (14). Its broad spectrum makes it an important agent for reducing the usage of carbapenem drugs, which are globally important last line antibiotics. Limiting carbapenem use is a critical element of antimicrobial stewardship and essential for preventing the spread of resistance (15). Treatment options for carbapenem resistant bacteria are often limited to poorly tolerated drugs (e.g. colistin or tigecycline) (16). Whilst TZP does possess in-vitro activity against ESBLs, the MERINO trial did not demonstrate non-inferiority of TZP to meropenem in treating patients with ESBL *E. coli* and *K. pneumoniae* blood stream infections (17). Carbapenems are therefore now recommended for this patient group (18).

In 2018, resistance to TZP occurred in 9.1% of invasive *E. coli* isolates in the UK (19). This can be caused by the production of carbapenemase enzymes (20), multiple β-lactamases (21) or ESBLs in combination with increased efflux or porin loss (22), which also provide resistance to 3GCs. Recently, a phenotype of resistance to TZP with susceptibility to 3GCs (TZP-R/3GC-S) emerged in *E. coli* and *Klebsiella pneumoniae,* indicating the possibility of alternative resistance mechanisms. The major cause of this phenotype is the hyperproduction of class A or D β-lactamases such as TEM-1(23, 24). Increased production of β-lactamase overcomes the inhibitive effect of tazobactam, ostensibly through saturation of the inhibitor, allowing the excess enzyme to hydrolyse and degrade piperacillin (25). β-lactamase hyperproduction can occur via increased gene expression modulated by a stronger promoter (26), or an increase in gene copy number mediated by insertion sequences (27, 28) or plasmids (24). Other mechanisms have also been identified, including expression of OXA-1 (25), inhibitor resistant enzymes such as *b/a*_TEM-33 (11)_, and a single nucleotide polymorphism (SNP) at position S133G in *b/a*_CTX-M-15_ found *in vitro* via random mutagenesis/error prone PCR but not yet found in clinical isolates (29).

Routine blood culture surveillance identified the occurrence of this phenotype in *E. coli* at the Royal Liverpool University Hospital (RLUH), Liverpool, UK, between 2014 and 2017. We sought to identify the diversity of *E. coli* strains and distribution of known mechanisms of TZP. We compared our collection to the findings of a UK-wide collection of invasive *E. coli* isolates (*n* = 1509) with publicly available phenotypic and genotypic data. This data set included the TZP-R/3GC-S phenotype as well as a piperacillin/tazobactam and third generation cephalosporins susceptible (TZP-S/3GC-S) phenotype

## Methods

### Study setting

The RLUH is a city centre located hospital in Liverpool, UK, providing secondary and tertiary care, with a catchment area of >2 million people in Merseyside, Cheshire, North Wales, and the Isle of Man. In 2019 the hospital recorded over 587,000 outpatient appointments and 95,000 daily inpatients.

### Ethics statement

The study utilised bacterial isolates collected by the RLUH for standard diagnostic purposes. All isolates were anonymised and de-linked from patient data. As no human samples or patient data were utilised in the study, ethical approval was not required. This was confirmed using the online NHS REC review tool http://www.hra-decisiontools.org.uk/ethics/.

### Surveillance data & Isolate collection

Blood stream bacterial pathogens were isolated using the BacTAlert 3D blood culture system (bioMérieux, France) and identified to a species level using MALDI-TOF (Bruker, US). Antimicrobial susceptibility testing (AST) was carried out using disk diffusion-based testing according to the British Society of Antimicrobial Chemotherapy guidelines (30) between 2014 and August 7^th^ 2017, after which these were replaced by the European Committee for Antimicrobial Susceptibility Testing (EUCAST) guidelines (31). In 2014 ceftazidime was used as the indicator 3GC, which was changed to cefpodoxime between 2015 and 2017. Isolate details and AST results were recorded in the Laboratory Information System (Telepath, CSC, US). All isolates were retained in glycerol stocks at −80°C in the RLUH Biobank. Data for the study was extracted into a database, including susceptibility data for ampicillin, cefpodoxime/ceftazidime, TZP, meropenem, ertapenem, cefoxitin, ciprofloxacin, gentamycin, amikacin, amoxicillin/clavulanic acid, tigecycline, and chloramphenicol. The data was used to estimate the proportion of *E. coli* isolates per year with TZP resistance, with and without associated 3GC resistance. In cases where multiple isolates were obtained from a single infectious episode, only the first isolate was included for further investigation and sequencing, to avoid duplication. Isolates that were TZP-R/3GC-S were retrieved from the Biobank and resurrected from glycerol stocks using Luria-Bertani agar (Oxoid, UK) and incubated at 37°C for 18 hours.

### Antimicrobial Susceptibility

Minimum inhibitory concentrations (MIC) for the isolates were obtained using the E-TEST method (Biomerieux, France) (32) according to EUCAST guidelines. (33) MICs were determined for TZP and the 3GC ceftriaxone (CTX).

### DNA extraction and sequencing

Genomic DNA was extracted using the PureGene Yeast/Bacteria Kit (Qiagen, Germany), following the manufacturer’s instructions for Gram-negative bacteria. Genome sequencing of 65 isolates was performed by MicrobesNG (http://www.microbesng.uk), using 2 × 250 bp short-read sequencing on the IIIumina MiSeq (IIIumina, US) (Table S1).

### Genome analysis, sequence typing and AMR gene prediction

All genomes were *de novo* assembled and annotated using SPAdes version 3.7 (34), and Prokka 1.11 (35), respectively, by MicrobesNG, in addition to providing the trimmed and quality filtered sequencing reads. The presence and copy number of AMR genes was determined using ARIBA (36), with the SRST2 database (37). *In silico* multi locus sequence typing (MLST), and plasmid replicon typing were carried out using ARIBA and the MLSTFinder (38) and PlasmidFinder (39) databases, respectively. β-lactamase promoters were identified by constructing databases with promoter sequences for *b/a*_TEM-1_ (26) and screening using ARIBA. Copy numbers were estimated by dividing the sequencing coverage of β-lactamase genes by the coverage of the chromosomal single copy gene *ampH.*

### Phylogenetic analysis of study isolates

A pan-genome analysis of all sequences was generated using Roary (40), and the core gene alignment was used as input for snp-sites (41) to extract ACGT-only SNPs (-c option). A maximum likelihood tree was produced using iqtree (42), with the general time reversible (GTR) model and gamma correction using ASC ascertainment bias correction (ASC) for SNPs-only alignments (-m GTR+G+ASC) and 1000 bootstrap replicates (-bb 1000). Phylogenetic trees were annotated using the Interactive Tree of Life (43) (https://itol.embl.de/). Core genome trees for sequence types ST131 and ST73 were generated by mapping the reads against the reference chromosomes of *E. coli* strains EC958 (HG941718.1) and CFT073 (AE014075.1), respectively, using snippy (https://github.com/tseemann/snippy). Recombination blocks were removed with Gubbins (44), and extraction of SNPs-only of the recombination-free alignment, and tree calculation, were performed as described above, using SNP-sites and IQ-TREE.

To investigate the relation of the study isolates to the whole UK hospital *E. coli* population, the sequences from a large UK-wide comparative analysis were included (PRJEB4681, (45)). These sequences included 1094 isolates submitted to the UK wide Bacteraemia Resistance Surveillance Programme (www.bsacsurv.org) between 2001-2011 by 11 hospitals across England, and 415 isolates provided by the Cambridge University Hospitals NHS Foundation Trust, Cambridge.

A core gene alignment and phylogenetic tree were constructed. Isolates from the UK-wide collection with the same phenotype of TZP resistance/3GC susceptibility (defined as susceptibility to both ceftazidime and cefotaxime, or either compound if only one was tested) were identified from the phenotypic AMR data (45), and highlighted alongside study isolates.

### Data availability

Raw read data and assemblies were submitted under BioProject ID PRJNA644114. Detailed per-strain information on accession numbers, resistance profiles, resistance gene predictions and sequence types (STs) are given in Table S1.

## Results

### Isolate collection and antimicrobial susceptibility testing

The RLUH recorded 1472 BSI *E. coli* isolates between 2014 and 2017 and antimicrobial susceptibility testing showed 172 isolates (11.8%) were resistant to TZP (Fig.S1). The proportion of *E. coli* resistant to TZP declined between 2014 (21%) and 2017 (9%, Fig. 1C). Of the 1258 TZP-susceptible isolates, the majority (1129, 89.7%) were susceptible to 3GC, while 129 (10.3%) were 3GC non-susceptible. In contrast, 86/172 (50%) TZP-resistant isolates were non-susceptible and 86/172 (50%) were susceptible to 3GC (Fig.1A).

**Fig. 1.**
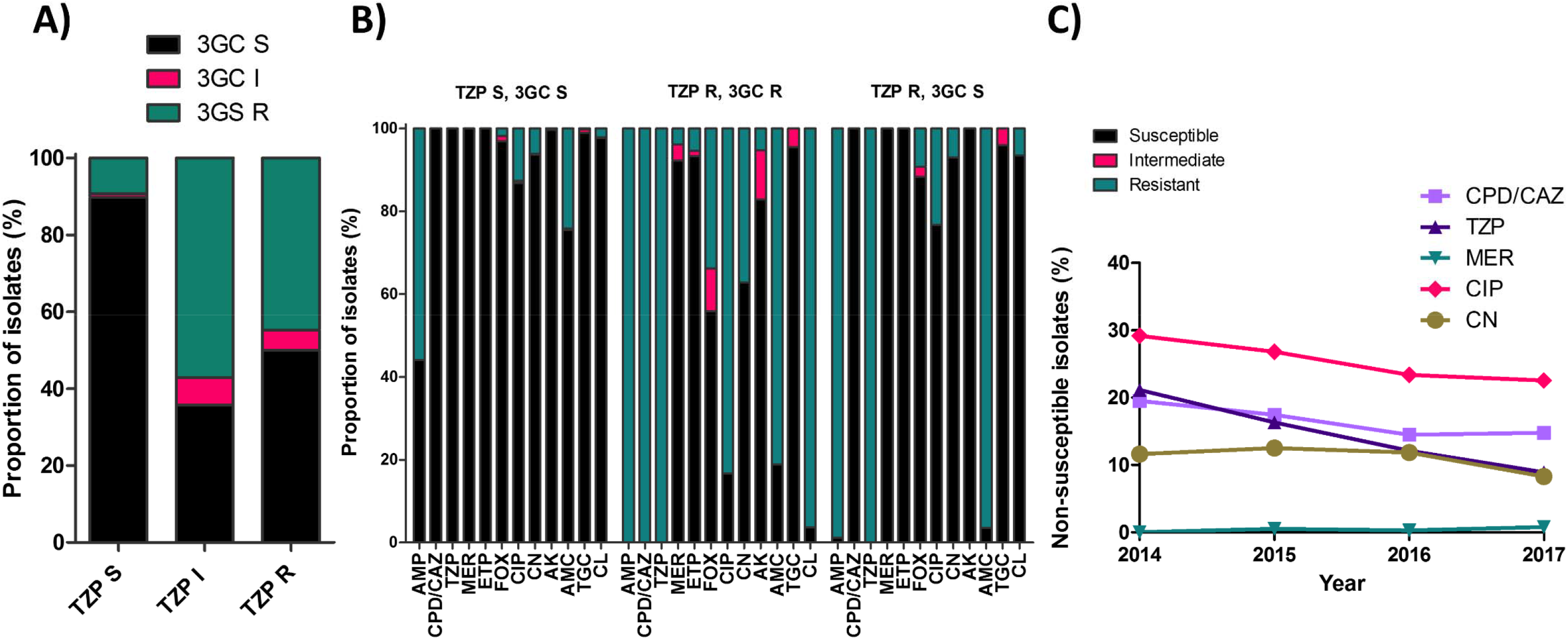
(A) Proportion of TZP susceptible (TZP S), intermediate (TZP I) and resistant (TZP R) isolates that are third generation cephalosporin susceptible (3GC S), intermediate (3GC I) and resistant (3GC R). (B) Antimicrobial susceptibilities of E. coli isolates from the RLUH, grouped by their susceptibility to piperacillin/tazobactam (TZP) and third generation cephalosporins (3GC). Susceptibility data is shown for isolates that are TZP susceptible and 3GC susceptible (TZP S, 3GC R). TZP resistant and 3GC resistant (TZP R, 3GC R), and TZP resistant and 3GC susceptible (TZP R, 3GC S), for the antibiotics ampicillin (AMP), cefpodoxime/ceftazidime (CPD/CAZ), piperacillin/tazobactam (TZP), meropenem (MER), ertapenem (ETP), cefoxitin (FOX), ciprofloxacin (CIP), gentamycin (CN), amikacin (AK), amoxicillin/clavulanic acid (AMC), tigecycline (TGC), and chloramphenicol (CL). (C) Trends in non-susceptibility to CPD/CAZ, TZP, MER, CIP and CN between 2014 and 2017 at RLUH.

Resistance to carbapenems was only seen in the TZP-resistant/3GC-resistant isolates, with 3.9% resistant to meropenem and 5.3% to ertapenem. A higher proportion of the TZP-R/3GC-S isolates were resistant to amoxicillin/clavulanic acid in comparison with TZP-resistant/3GC-resistant isolates (96.4% vs 81.1%) (Fig.1B). Overall, aside from the penicillin class antibiotics, the TZP-R/3GC-S phenotype had high incidence of susceptibility towards the antimicrobials tested.

Of the 86 isolates with the TZP-R/3GC-S phenotype, 14 had been derived from repeated sampling of long-term patients and were excluded, resulting in 72 isolates derived from unique patients. These isolates were reduced to 66 after excluding TZP MICs under the EUCAST breakpoint for susceptibility. A further isolate was considered a contaminant *(Staphylococcus aureus)* based on colony morphology, which was confirmed by 16S PCR. After whole genome sequencing, two of the 65 isolates were removed as they contained more than one *E. coli* genome, either due to mixed infections or contamination (assembly sizes were 9602556bp and 9552068bp, respectively), leaving 63 isolates for further analysis.

The MICs of TZP as assessed by the E-TEST ranged from 12 to 256 mg/L. Fifty-eight isolates had MICs over the EUCAST breakpoint for resistance (16 mg/L), and five had intermediately resistance (MIC 12mg/L). The CTX MICs ranged between 0.016 and 0.25 mg/L, all below the breakpoint for resistance (2mg/L), confirming the TZP-R/3GC-S phenotype.

### Resistance and plasmid profile of TZP-resistant/3GC-susceptible population

The AMR genotypes (Fig.S2) correlated well with the phenotypic data obtained by disk testing, with most isolates susceptible to ciprofloxacin and gentamicin. The 58 TZP-R/3GC-S isolates harboured a variety of β-lactamase genes, including TEM-type (n=44; *b/a*_TEM-1_ [41], *b/a*_TEM-33_ [2], *b/a*_TEM-148_ [1]), *b/a*_SHV-1_(n=9), *b/a*_CTX-M-15_ (n=4) and *b/a*_OXA-1_ (n=3)).The presence of β-lactamase genes correlated with resistance to ampicillin and TZP, whilst resistance to ciprofloxacin in 17/58 isolates (29%) was accounted for by *gyrA* mutations D87N (10/17, 59%) and S83L (12/17, 71%),and *parC* S80I mutation (10/17, 59%). Aminoglycoside resistance was explained by the *O-adenylyltransferases aadA* (6/6, 100%), in combination with the genes *aac(3)-IIa* or *aadB* (3/6, 50%). Additionally, all isolates carried the chromosomal *b/a*_AmpC1_ which is constitutively expressed at a low level (46), and 51/58 of the isolates carried *b/a*_AmpC2_. A single isolate (169961) had a coding mutation in a penicillin binding protein, with an A37T mutation in *mrdA* encoding penicillin binding protein 2, in combination with the inhibitor resistant *b/a*_TEM-33_ and strong *Pa/Pb* promoter.

Replicons usually associated with large resistance plasmids, such as lncFIA and lncFIB, lncFIA and lncFIIA, were detected in 19% of the study isolates (Fig.2), reflecting the low proportion of isolates with multiple resistance genes and the unusual resistance profile characteristic of the TZP-R/3GC-S phenotype.

**Fig. 2.**
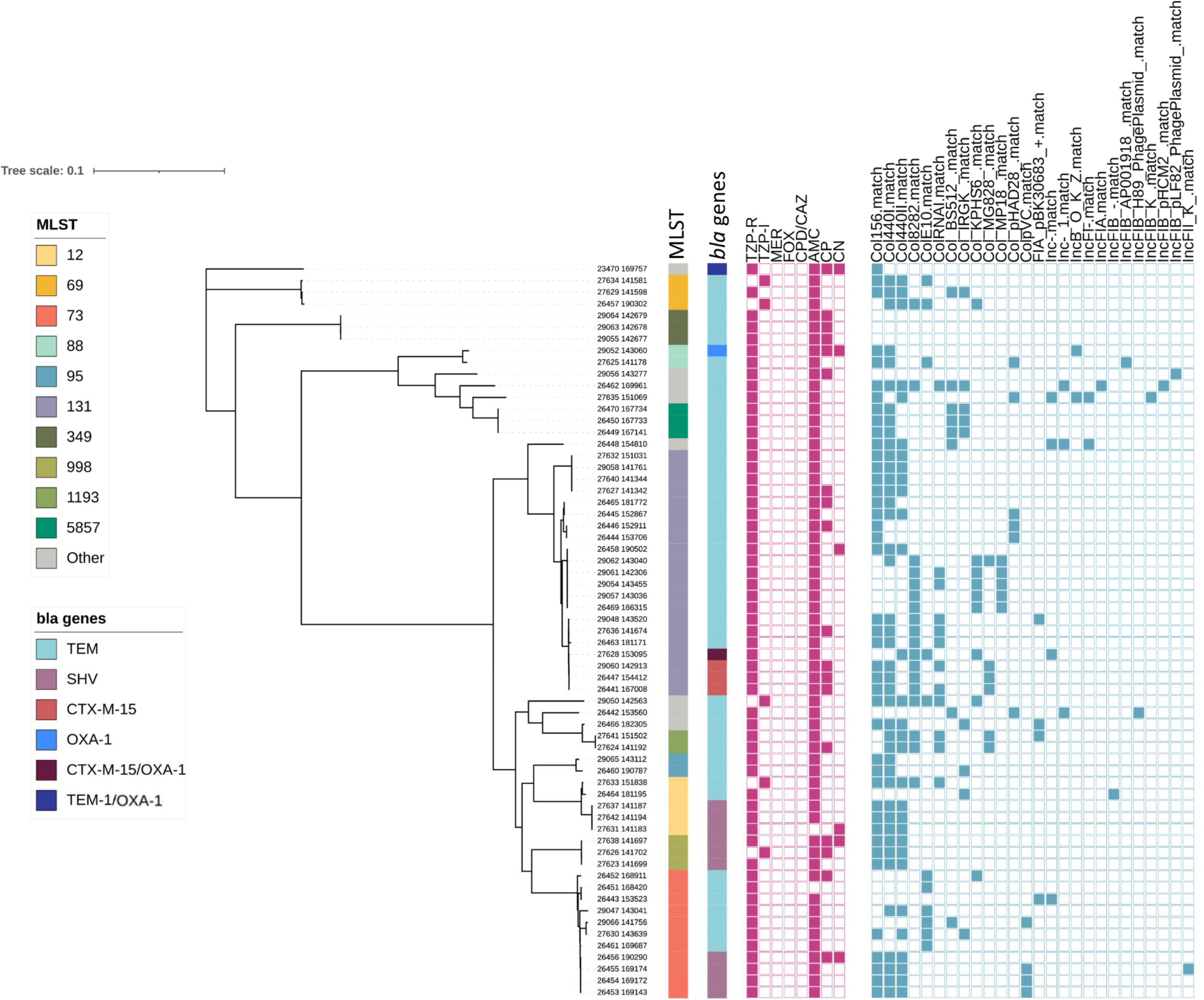
Maximum likelihood phylogeny of the study isolates from RLUH. The colour strips, from left to right, show the MLST classification (MLST), β-lactamase gene carriage (*b/a* genes). The heat maps show phenotypic resistance to piperacillin/tazobactam (TZP), meropenem (MER), cefoxitin (FOX), cefpodoxime/ceftazidime (CPD/CAZ), ampicillin (AMP), ciprofloxacin (CP), and gentamycin (CN), and the plasmid replicon repertoire.

### Population structure of the TZP-resistant/3GC-susceptible population within the nationwide context

Phylogenetic analysis revealed the TZP-R/3GC-S phenotype occurred in a diverse number of sequence types (Fig.2). The 58 TZP-R/3GC-S isolates represented 16 STs. The most representative were ST131 (36.2%), ST73 (19%) and ST12 (6.9%). The TZP-R/ 3GC-S phenotype in the UK-wide collection was similarly diverse to the RLUH collection with ST131 (22.1%) the most represented, followed by ST73 (16.2%) and ST95 (14.7%, Fig.3). This diversity was also reflected in the TZP-S/3GC-S phenotype; ST73 (16.8%), ST131 (14.3%) and ST95 (10.6%). When placing the RLUH isolates into the phylogenetic context of the UK-wide bloodstream isolates collected from 2001 to 2011, it was apparent that they reflected the overall *E. coli* population structure (Fig.4). This indicates that the TZP-R/ 3GC-S phenotype is not driven by a clonal outbreak within this single hospital setting, but rather by multiple acquisitions of resistance mechanisms in the circulating population of hospital strains. The AMR gene profile of the RLUH isolates varied between STs (Fig.S3), with ST131 carrying more resistance genes than the other major STs, as previously reported (47). To get a higher-resolution insight into the within-ST diversity of the isolates, we calculated core genome trees of the main STs by mapping the reads against selected reference genomes and extracting the conserved, non-recombinant SNPs. The acquisition of the phenotype was not a single event even in these closely related organisms, as it occurred on several occasions for both main sequence types, with no (ST73; Fig. S4) or very few (ST131; Fig. S5) isolates closely related, which may indicate within-hospital transmission.

**Fig. 3.**
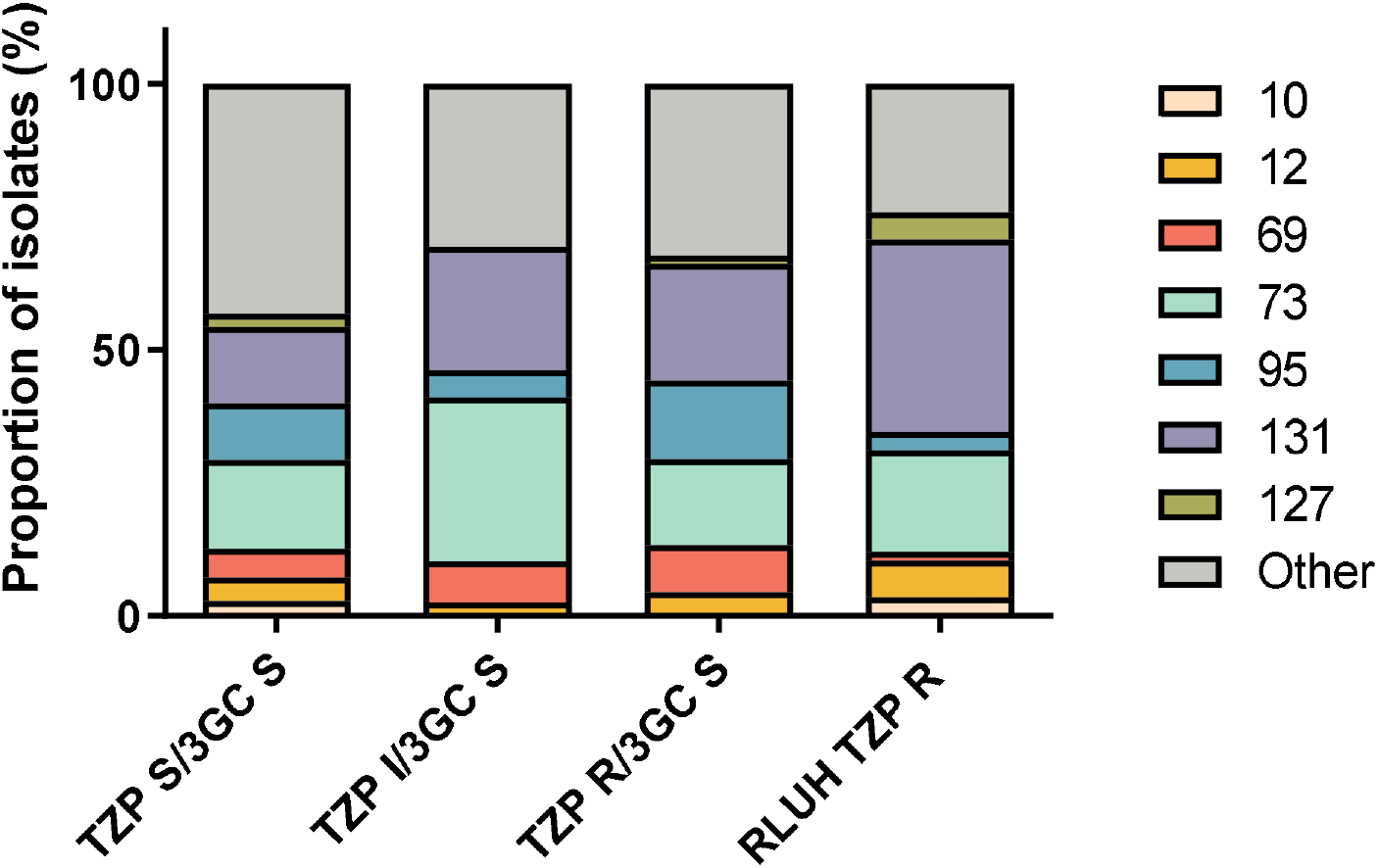
Bar chart showing the proportion of isolates belonging to common sequence types in the RLUH study isolates in comparison with those in the collection of 1509 isolates taken from a UK wide study (45).

**Fig. 4.**
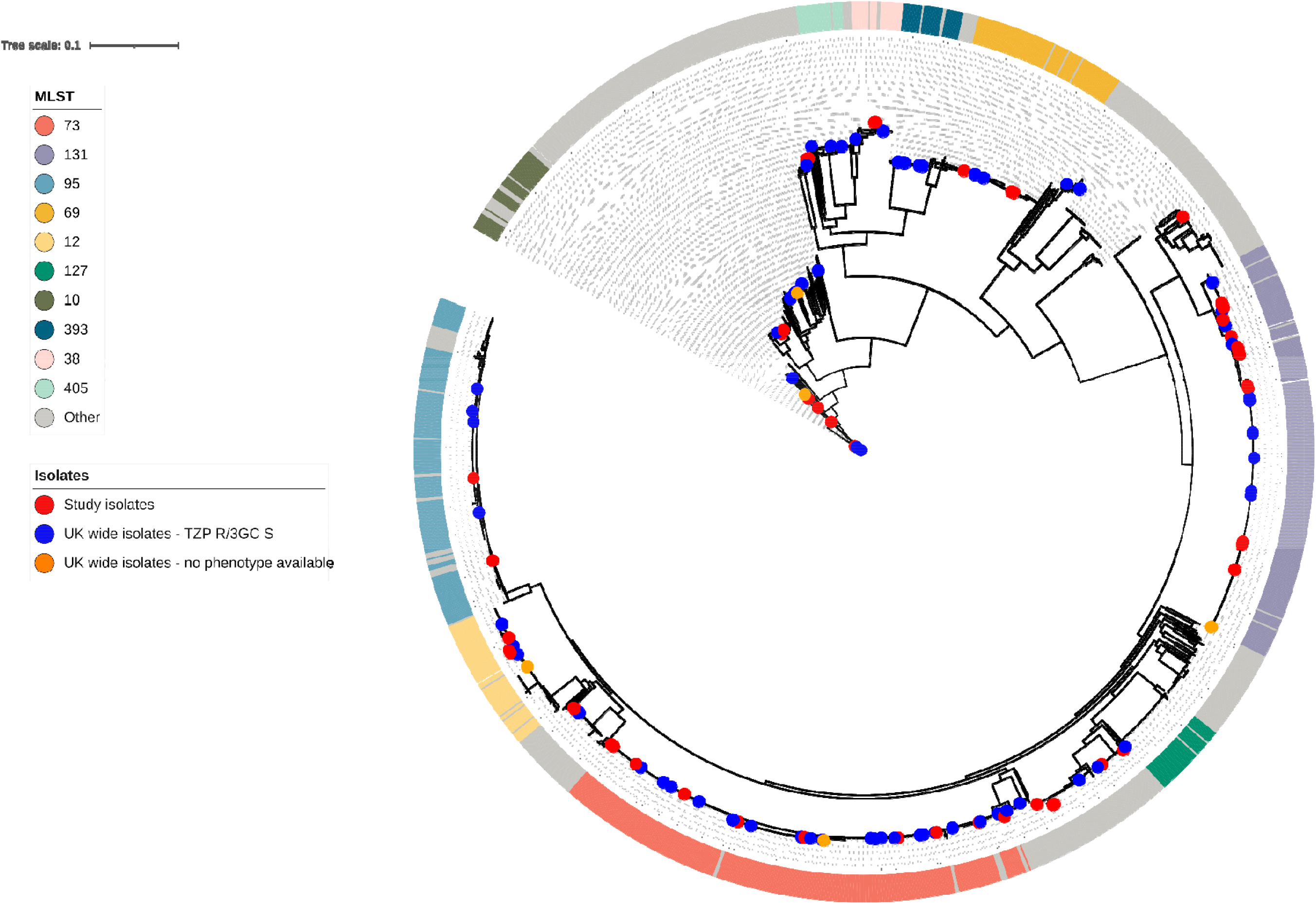
Circular Maximum Likelihood core genome phylogenetic tree of the 68 study isolates in combination with 1509 UK wide study isolates. The ring indicates the ten most commonly encountered STs. Dots at the terminus of branches indicate study isolates, UK wide isolates with the TZP resistant/3GC susceptible phenotype (TZP-R/3GC -S) or isolates from the UK wide collection missing sufficient phenotypic data to assign an accurate AMR phenotype.

### Varied putative genetic determinants of the TZP-resistant/3GC-susceptible phenotype

We sought to identify previously published putative resistance mechanisms associated with the TZP-R/3GC-S phenotype in the 58 isolates from RLUH (Fig. 5A). No carbapenemase genes were predicted to be present, although four ST131 isolates harboured the ESBL *b/a*_CTX-M-15_ gene, normally associated with 3GC resistance. However, three isolates carried a SNP resulting in the non-synonymous amino acid change from serine to glycine at position 133. This amino acid change is reported to result in a non-ESBL phenotype with increased TZP resistance. However the S133G mutation was only identified through random mutagenesis/error prone PCR *in vitro* (29). In the remaining isolate with *b/a*_CTX-M-15_ the promoter sequence was deleted and therefore presumably not expressed (Fig, S6). However, the isolate also carried *b/a*_OXA-1_. In all three isolates carrying *b/a*_OXA-1_, it was either the sole β-lactamase or it was carried with a second β-lactamase. Of the isolates with *b/a*_TEM-1_, 25 had the weak *P3* promoter, four had the strong promoter *P4* and 12 contained the strong, overlapping promoter *Pa/Pb.* The *P4* and *Pa/Pb* promoter have previously been linked to hyperproduction of TEM-1 (26). The TZP-R/3GC-S phenotype has previously been associated with increases in the copy number of *b/a*_TEM-1_ via gene amplification, resulting in hyperproduction of the TEM-1 enzyme (23, 27). The copy numbers of *b/a*_TEM-1_, as estimated by sequencing coverage, for those isolates within the RLUH collection with a weak *P3* promoter varied between 3 and 186 copies, and a mean of 44 copies (Fig. 5B).

**Fig. 5.**
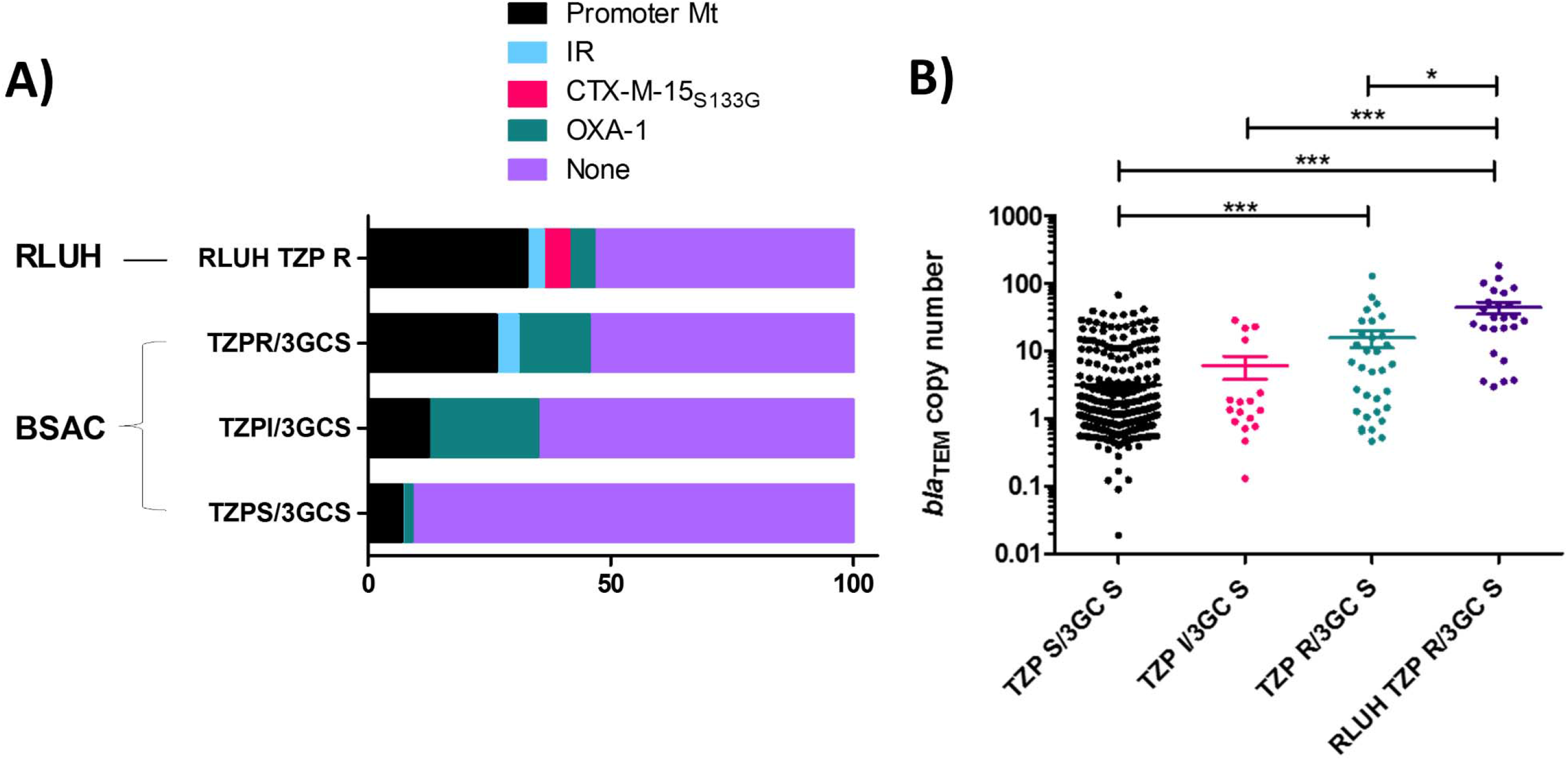
The proportion of isolates of each phenotype identified in the RLUH and BSAC collections with identifiable putative TZP resistance mechanisms (IR; inhibitor resistance) (A), and the copy number of *b/a*_TEM-1_ genes found in isolates belonging to each phenotype (B), with significance determined by Dunn’s Multiple Comparison Test.

We identified inhibitor resistant β-lactamases (3; 4%), *b/a*_OXA-1_ (10, 14.7%) and *b/a*_TEM-1_ promoter region mutations (18; 26%) in the 68 TZP-R/3GC-S isolates from the UK wide collection (Fig. 5A). However, we also identified these mechanisms, although at a lower incidence (inhibitor resistant β-lactamase; 2 [0.2%], *b/a*_OXA-1_ 24 [1.8%] promoter region mutations; 89 [7%]), in the TZP-S/3GC-S phenotype in the same collection. In total a putative mechanism was found in 27 of the 58 isolates.

The copy number of *b/a*_TEM-1_ was also elevated in both the TZP-R/3GC-S (min-max of 0.5 and 129 copies, mean 16 copies) and TZP-S/3GC-S (min-max of 0.02 and 68 copies, mean 3 copies) from the UK wide collection (Fig. 5B). Despite this, there was a significant difference in copy number between the TZP-S/3GC-S vs UK wide TZP-R/3GC-S phenotypes (P value < 0.001; Dunn’s Multiple Comparison Test) and the TZP-S/3GC-S vs TZP-R/3GC-S phenotypes from RLUH (P value < 0.001; Dunn’s Multiple Comparison Test). This indicates that although an increase in copy number of *b/a*_TEM-1_ may not be predictive of TZP-R/3GC-S, it is associated with the phenotype.

## Discussion

This phylogenetic analysis of the TZP-R/3GC-S phenotype in *E. coli* from RLUH demonstrates that this phenotype derives from repeated, multiple acquisition events. Our comparison of the RLUH isolates with a large UK-wide collection (45) shows that this is not unique to our study site, but broadly reflective of the phenotype from multiple sites across the UK. As the TZP-R/3GC-S phenotype also reflects the overall UK population structure *E. coli* bacteraemia isolates, this is suggestive of the impact of repeated or sustained antimicrobial pressure, rather than fixation in a certain lineage and subsequent spread. The phenotype was encountered in the typically drug resistant ST131 (48), and the often highly virulent but drug susceptible ST73 (49), reflecting the overall dominance of these STs, and was not associated with an overall increase in carriage of genes conferring resistance to other classes of antibiotics. We also were unable to identify the presence of a common plasmid replicon, further highlighting the diversity of the phenotype.

Strategies to increase the effectiveness of TZP include increasing dosage, which in one study increased the coverage of TZP from 83.2% to 93% of bacterial blood stream pathogens (50). Increasing the concentration of tazobactam alongside a fixed dose of piperacillin has also rescued TZP effectiveness against TEM-1 hyperproducers in a neutropenic mouse model (25), and could be a viable strategy to protect its future effectiveness. It is worth noting observational clinical data (51), and *in vivo* experimental data (52), suggesting TZP may be effective against some organisms with *in vitro* phenotypic resistance to TZP.

The rapid identification of the TZP-R/3GC-S phenotype would enable de-escalation from TZP to a 3GC (53), both reducing the likelihood of treatment failure, and preventing overuse of carbapenems, which is key for antimicrobial stewardship (54). The isolates were also mostly susceptible to ciprofloxacin, gentamicin and amikacin, providing further de-escalation opportunities. Recent work on methicillin resistant *Staphylococcus aureus* has described frequent collateral sensitivity to narrow spectrum penicillin/inhibitor combinations, highlighting that targeted de-escalation rather than escalation can be possible when treating organisms highly resistant to first line drugs (55). Molecular diagnostics and whole genome sequencing can be used to rapidly detect AMR genes to predict AMR phenotype (56, 57). However, it is essential to match the phenotypic resistance to the genotypic mechanisms. The majority of TZP-R/3GC-S isolates in this study hyperproduced the class A β-lactamase enzymes *b/a*_TEM-1_, which can hydrolyse piperacillin but not 3GCs, and is inhibited by tazobactam. Hyperproduction can occur via gene amplification, in which tandem repeats of AMR genes are generated, for example via the 1S26 mediated amplification of pseudo-compound transposons (27, 58), or the transfer of β-lactamase genes to high copy AMR plasmids (24). A number of the isolates were lacking a detectable increase in gene copy number, but had a potential route to hyperproduction via a strong promoter of *b/a*_TEM-1_ (26).

We also detected *b/a*_TEM-33_, encoding an inhibitor resistant variant of TEM-1B (59), and *b/a*_OXA-1_, either as the only β-lactamase or in combination with *b/a*_TEM-1_. OXA-1 is poorly inhibited by tazobactam (60) but has been associated with the TZP-R/3GC-S phenotype (61), while a recent UK study identified *b/a*_OXA-1_ as a major contributor to TZP resistance amongst ESBL *E. coli* (61). However, the carriage of *b/a*_OXA-1_ does not always confer resistance to TZP, which appears to depend on the genetic background of the strain. The risk ratio of *b/a*_OXA-1_ being associated with TZP resistance in ESBL *E. coli* is higher in ST131 strains (12.1) compared with ESBL *E. coli* as a whole (6.49) (61). One isolate carrying the OXA-1 β-lactamase gene, as well as *b/a*_CTX-M-15_ lacking a promoter, belonged to ST131.

Three out of four detected *b/a*_CTX-M-15_ encoded the S133G mutation, which increases TZP MIC ten-fold, whilst reducing the 3GC MIC by the same margin in a strain harbouring a random mutagenesis/error prone PCR derived *b/a*_CTX-M-15_(29). To our knowledge this is the first report of this *b/a*_CTX-M-15_ variant in clinical isolates. The S133G mutation in *b/a*_CTX-M-15_ was associated with 5% of TZP-R/3GC-S in our setting and only in ST131. The mutation of *b/a*_CTX-M_ genes to better hydrolyse mecillinam has been reported during urinary tract infections treatment (62), but not for TZP or other β-lactam/inhibitor combinations. The circulation of *b/a*_CTX-M_ variants that do not confer the ESBL phenotype but provide resistance to TZP, has implications for molecular testing for ESBL organisms (63), as it would misclassify the isolates as 3GC-resistant and lead to unnecessary use of carbapenems. There is thus a need for *in vitro* development of resistant mutations to uncover potential routes to resistance and improve AMR prediction. We found that 9 of 58 isolates, all without a putative resistant mechanism, harboured blaSHV-48. Hyperproduction of this enzyme has been shown to lead to the TZP-R/3GC-S phenotype in *Klebsiella pneumoniae* (64).

All the putative mechanisms of TZP-R/3GC-S found in the isolates from RLUH have been previously published and widely associated with this phenotype. However, we found evidence of *b/a*_TEM-1_ promoter region mutations, inhibitor resistance enzymes and increased *b/a*_TEM-1_ copy number in the TZP-S/3GC-S phenotype. The only putative mechanism which was not found in the TZP-S/3GC-S phenotype was the S133G mutation in *b/a*_CTX-M-15_, This mutation was not found in any of the TZP-R/3GC-S phenotype isolates from the UK wide collection, which may indicate low incidence or a localised emergence in our hospital. The diverse putative mechanisms of TZP-R/3GC-S and phenotype-genotype discordance, as seen in TZP-S/3GC-S, would compromise current molecular or genomic detection of this phenotype.

The main limitation of this study was that only TZP-R/3GC-S isolates from the RLUH were sequenced, and the relatively small population size. We utilised a large and UK-wide collection of isolates for comparison, which were similarly diverse and reflected the overall population structure (45).

This work highlights the phylogenetic diversity of the TZP-R/3GC-S phenotype in *E. coli* and the variety of the putative resistance mechanisms involved, including β-lactamase hyperproduction via gene amplification and promoter mutations, inhibitor resistant TEM-1 and CTX-M-15 variants. However, the presence of these mechanisms at a lower incidence with the TZP-S isolates highlights that a greater understanding of the evolution of TZP resistance and the resistance mechanisms of the TZP-R/3GC-S phenotype would be fundamental to improve the prediction of TZP-R/3GC-S *E. coli.* Until such time, phenotypic monitoring of this phenotype is essential to prevent treatment failure.

## Author contributions

TE, EH, JM, CMP and ATMH conceptualised the study. JvA, AH PR, CC, CMP, JM, and AH collated isolate metadata, and clinical antimicrobial susceptibility testing data. TE, EH, ERA, APR, LEC and ATMH contributed to the experimental design and data analysis. Bioinformatic analysis was carried out by TE and EH. TE, JvA, CTW, AJF, IB and ATMH carried out microbiological experiments. TE, EH and ATMH wrote the first draft of the manuscript. All authors reviewed and edited the final manuscript.

## Acknowledgments

We acknowledge the technical staff in the diagnostic microbiology laboratories in the RLUH and expert informatics support from the Pathogen Informatics team at the Wellcome Sanger Institute.

## Additional information

Correspondence and requests for materials should be addressed to TE or ATMH.

## Supplementary Figures

**Fig. S1.**
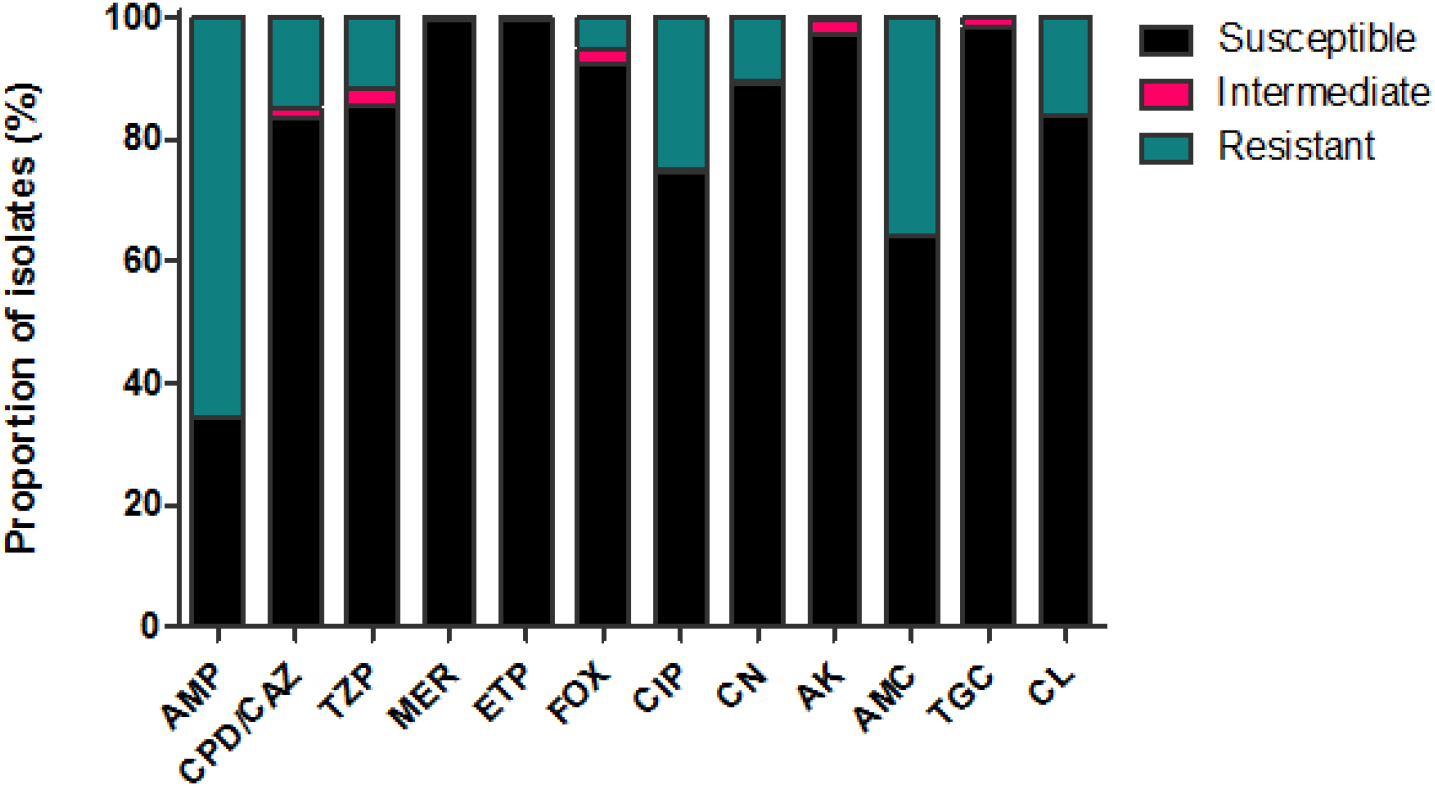
Proportion of the total E. coli isolated from blood stream infections between 2014 and 2017 at RLUH that were susceptible, intermediate or resistant to ampicillin (AMP), cefpodoxime/ceftazidime (CPD/CAZ), piperacillin/tazobactam (TZP), meropenem (MER), ertapenem (ETP), cefoxitin (FOX), ciprofloxacin (CIP), gentamycin (CN), amikacin (AK), amoxicillin/clavulanic acid (AMC), tigecycline (TGC), and chloramphenicol (CL).

**Fig. S2.**
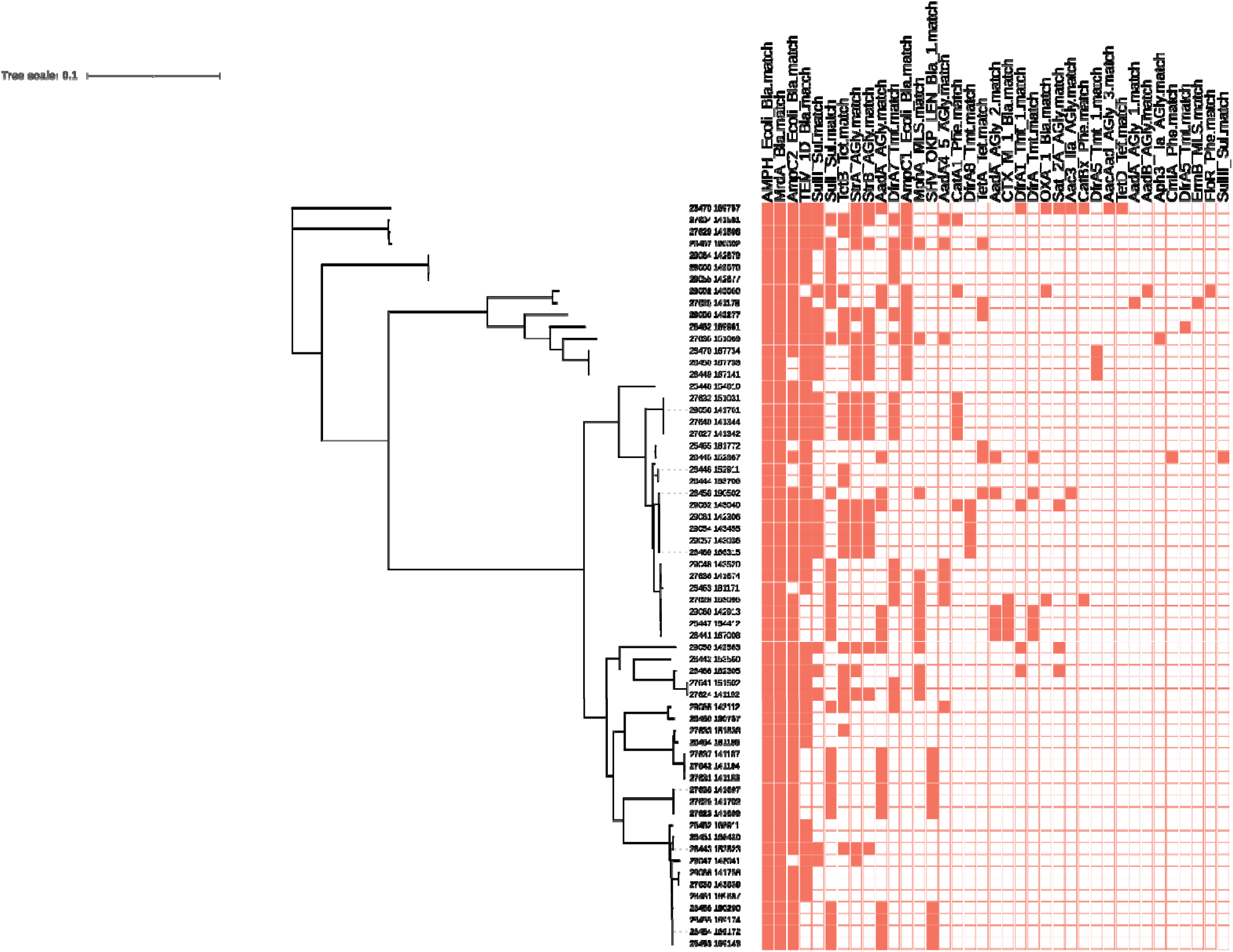
Maximum likelihood phylogeny of the study isolates from RLUH, with a heat map indicating the AMR gene repertoire.

**Fig. S3.**
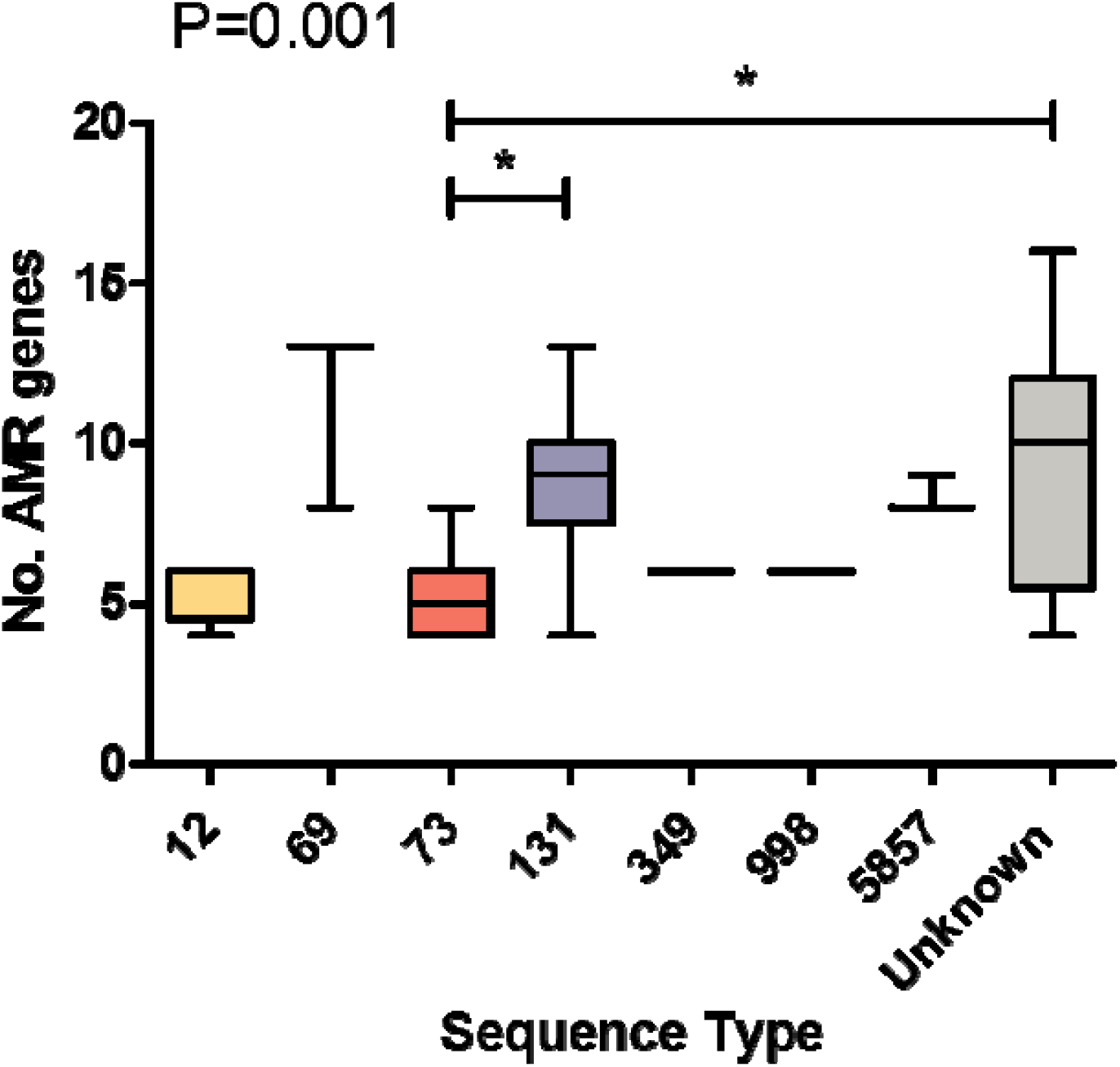
The number of AMR genes in isolates from the major sequence types encountered in the study. Whiskers show minimum and maximum values. Significance determined by Kruskal-Wallis test, * indicates a p value of <0.05.

**Fig. S4.**
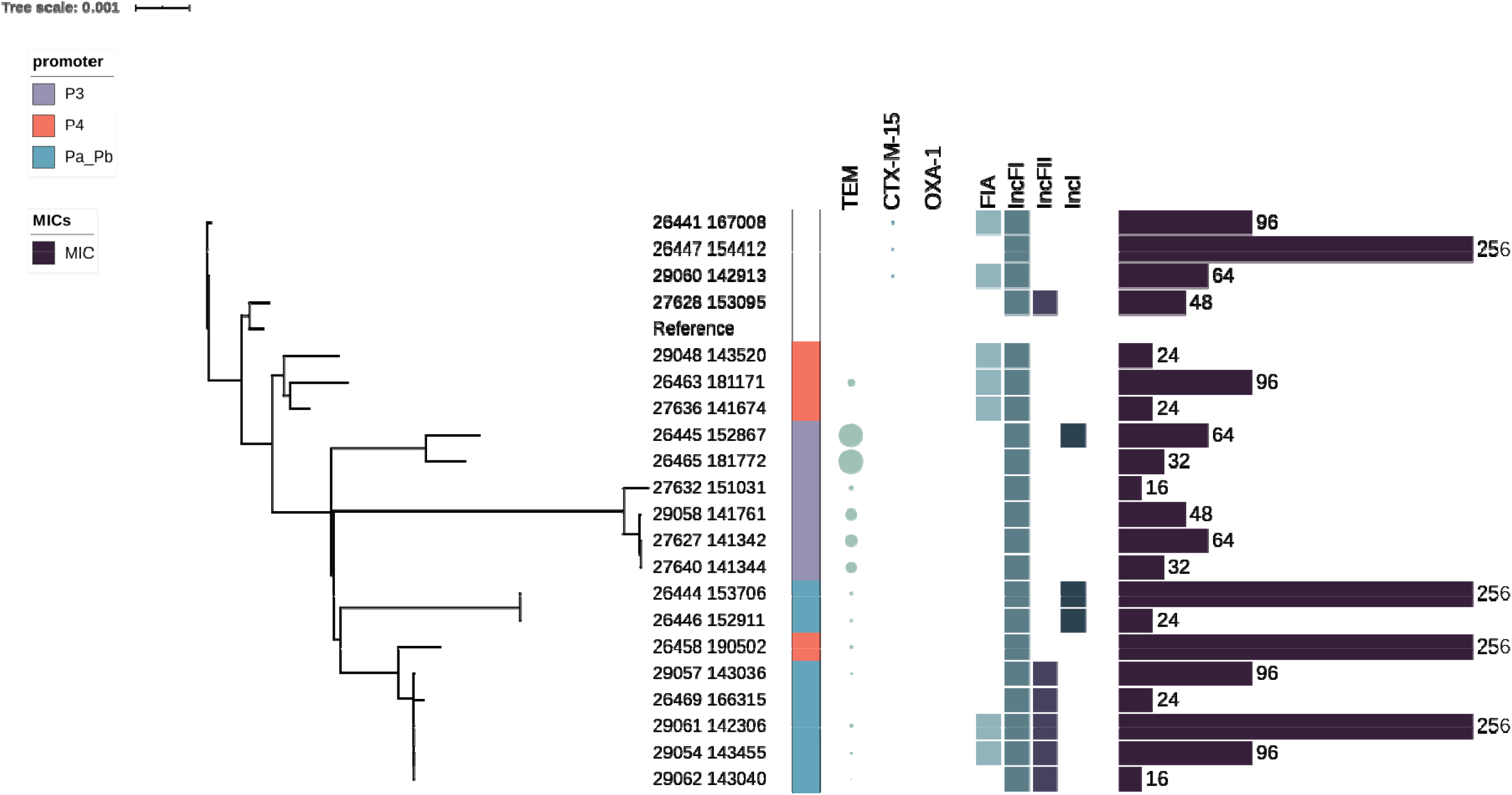
High resolution core genome – based phylogeny of TZP resistant/3GC susceptible ST131 isolates. Indicated are promoter types, β-lactamase copy numbers (size of circle represents relative copy number), plasmid replicons, and TZP MIC.

**Fig. S5.**
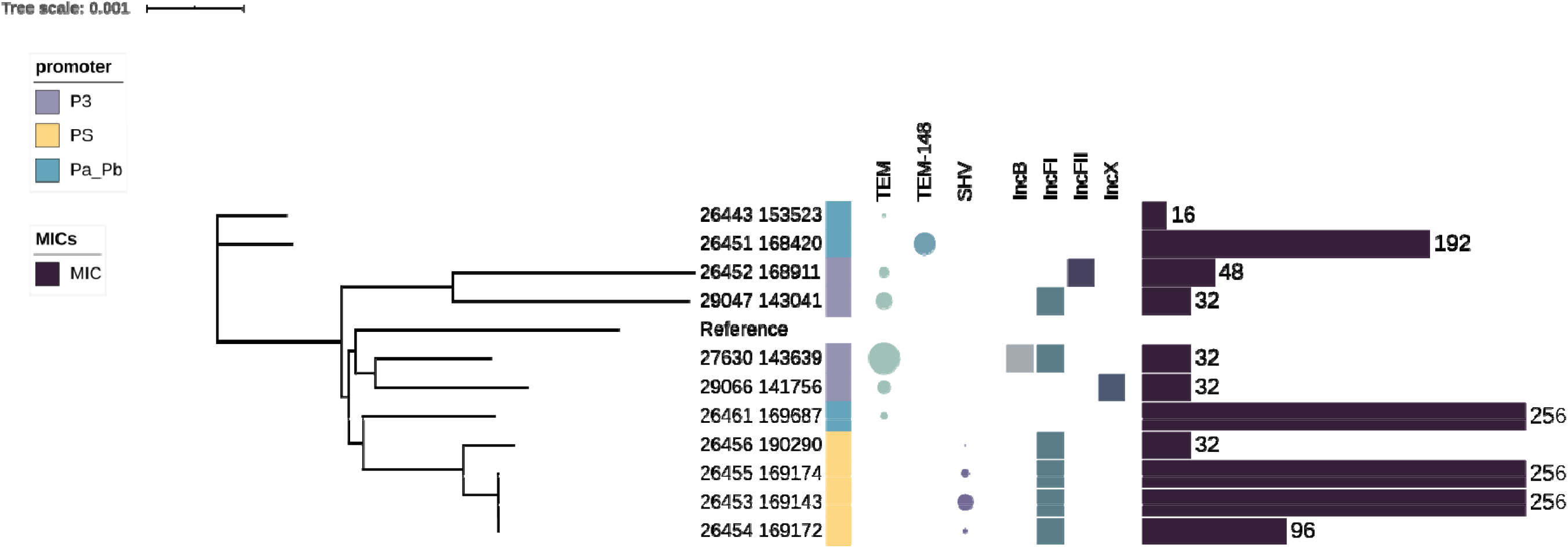
High resolution core genome – based phylogeny of TZP resistant/3GC susceptible ST73 isolates. Indicated are promoter types, β-lactamase copy numbers (size of circle represents relative copy number), plasmid replicons, and TZP MIC.

**Fig. S6:**
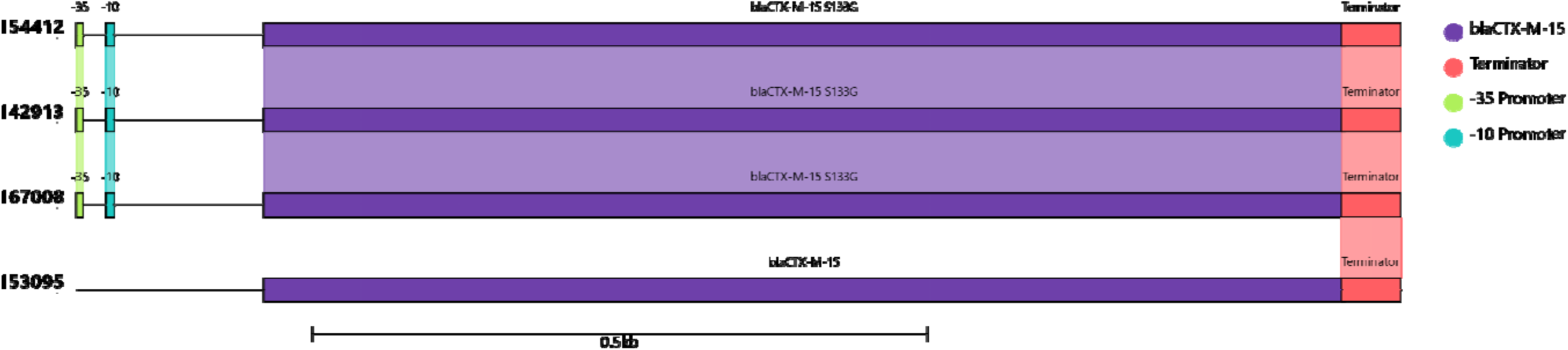
Comparison of the promoter region, gene and terminator of blaCTX-M-15 predicted to be present in four clinical isolates from the Royal Liverpool University Hospital collection. Isolates 154412, 142913 and 167008 all contained an intact promoter and terminator with a blaCTX-M-15 containing the S133G mutation, while isolate 153095 harboured a wild type blaCTX-M-15 but lacking the promoter. Shaded regions between isolates indicate 100% identity. Figure produced using clinker (65).

## References

1. Laupland KB. Incidence of bloodstream infection: a review of population-based studies. Clin. Microbiol. Infect. 2013;19(6):492–500.

2. Bonten M, Johnson JR, van den Biggelaar AHJ, Georgalis L, Geurtsen J, de Palacios PI, et al. Epidemiology of Escherichia coli Bacteremia: A Systematic Literature Review. Clin. Infect. Dis. 2020.

3. Public Health England. Annual epidemiological commentary: mandatory MRSA, MSSA and E coli bacteraemia and C difficile infection data. 2014.

4. World Health Organization. Global priority list of antibiotic-resistant bacteria to guide research, discovery, and development of new antibiotics. 2017.

5. Tacconelli E, Carrara E, Savoldi A, Harbarth S, Mendelson M, Monnet DL, et al. Discovery, research, and development of new antibiotics: the WHO priority list of antibiotic-resistant bacteria and tuberculosis. Lancet Infect. Dis. 2018;18(3):318–27.

6. World Health Organization. Landscape of diagnostics against antibacterial resistance, gaps and priorities. 2019.

7. Temkin E, Fallach N, Almagor J, Gladstone BP, Tacconelli E, Carmeli Y. Estimating the number of infections caused by antibiotic-resistant Escherichia coli and Klebsiella pneumoniae in 2014: a modelling study. Lancet Glob. health. 2018;6(9):e969–e79.

8. European Centres of Disease Control. Surveillance of antimicrobial resistance in Europe 2018 2018.

9. Tooke CL, Hinchliffe P, Bragginton EC, Colenso CK, Hirvonen VHA, Takebayashi Y, et al. β-Lactamases and β-Lactamase Inhibitors in the 21st Century. J Mol Biol. 2019;431(18):3472–500.

10. Gin A, Dilay L, Karlowsky JA, Walkty A, Rubinstein E, Zhanel GG. Piperacillin-tazobactam: a beta-lactam/beta-lactamase inhibitor combination. Expert review of anti-infective therapy. 2007;5(3):365–83.

11. Drawz SM, Bonomo RA. Three decades of beta-lactamase inhibitors. Clin. Microbiol. Rev. 2010;23(1):160–201.

12. Johnson DM, Biedenbach DJ, Jones RN. Potency and antimicrobial spectrum update for piperacillin/tazobactam (2000): emphasis on its activity against resistant organism populations and generally untested species causing community-acquired respiratory tract infections. Diagn. microbiol. Infect. Dis. 2002;43(1):49–60.

13. Kim M-K, Xuan D, Quintiliani R, Nightingale CH, Nicolau DP. Pharmacokinetic and pharmacodynamic profile of high dose extended interval piperacillin–tazobactam. J. Antimicrob. Chemother. 2001;48(2):259–67.

14. Barton GJ, Morecroft CW, Henney NC. A survey of antibiotic administration practices involving patients with sepsis in UK critical care units. Int J Clin Pharm. 2020;42(1):65–71.

15. Wilson APR. Sparing carbapenem usage. Journal of Antimicrobial Chemotherapy. 2017;72(9):2410–7.

16. Morrill HJ, Pogue JM, Kaye KS, LaPlante KL. Treatment Options for Carbapenem-Resistant Enterobacteriaceae Infections. Open Forum Infect. Dis. 2015;2(2):ofv050.

17. Harris PNA, Tambyah PA, Lye DC, Mo Y, Lee TH, Yilmaz M, et al. Effect of Piperacillin-Tazobactam vs Meropenem on 30-Day Mortality for Patients With E coli or Klebsiella pneumoniae Bloodstream Infection and Ceftriaxone Resistance: A Randomized Clinical Trial. Jama. 2018;320(10):984–94.

18. Gutiérrez-Gutiérrez B, Rodríguez-Baño J. Current options for the treatment of infections due to extended-spectrum beta-lactamase-producing Enterobacteriaceae in different groups of patients. Clin. Microbiol. Infect. 2019;25(8):932–42.

19. Public Health England. Laboratory surveillance of Escherichia coli bacteraemia in England, Wales and Northern Ireland. Health Protection Report. 2018;13(37).

20. van Duin D, Bonomo RA. Ceftazidime/Avibactam and Ceftolozane/Tazobactam: Second-generation β-Lactam/β-Lactamase Inhibitor Combinations. Clin. infect. Dis. 2016;63(2):234–41.

21. Lee J, Oh CE, Choi EH, Lee HJ. The impact of the increased use of piperacillin/tazobactam on the selection of antibiotic resistance among invasive Escherichia coli and Klebsiella pneumoniae isolates. Int J Infect Dis. 2013;17(8):e638–43.

22. Suzuki Y, Sato T, Fukushima Y, Nakajima C, Suzuki Y, Takahashi S, et al. Contribution of beta-lactamase and efflux pump overproduction to tazobactam-piperacillin resistance in clinical isolates of Escherichia coli. Int. J. Antimicrob. Agents. 2020:105919.

23. Schechter LM, Creely DP, Garner CD, Shortridge D, Nguyen H, Chen L, et al. Extensive Gene Amplification as a Mechanism for Piperacillin-Tazobactam Resistance in Escherichia coli. mBio. 2018;9(2).

24. Rodríguez-Villodres Á, Gil-Marqués ML, Álvarez-Marín R, Bonnin RA, Pachón-Ibáñez ME, Aguilar-Guisado M, et al. Extended-spectrum resistance to β-lactams/β-lactamase inhibitors (ESRI) evolved from low-level resistant Escherichia coli. J. Antimicrob. Chemother. 2019;75(1):77–85.

25. Abdelraouf K, Chavda KD, Satlin MJ, Jenkins SG, Kreiswirth BN, Nicolau DP. Piperacillin-Tazobactam-Resistant/Third-Generation Cephalosporin-Susceptible Escherichia coli and Klebsiella pneumoniae Isolates: Resistance Mechanisms and In vitro-In vivo Discordance. Int. J. Antimicrob. Agents. 2020;55(3):105885.

26. Zhou K, Tao Y, Han L, Ni Y, Sun J. Piperacillin-Tazobactam (TZP) Resistance in Escherichia coli Due to Hyperproduction of TEM-1 β-Lactamase Mediated by the Promoter Pa/Pb. Front Microbiol. 2019;10:833-.

27. Hubbard ATM, Mason J, Roberts P, Parry CM, Corless C, van Aartsen J, et al. Piperacillin/tazobactam resistance in a clinical isolate of Escherichia coli due to IS26-mediated amplification of *b/a*_TEM-1B_. Nat Comms. 2020;4915

28. Hansen KH, Andreasen MR, Pedersen MS, Westh H, Jelsbak L, Schønning K. Resistance to piperacillin/tazobactam in Escherichia coli resulting from extensive IS 26-associated gene amplification of bla TEM-1. J. Antimicrob. Chemother. 2019;74(11):3179–83.

29. Rosenkilde CEH, Munck C, Porse A, Linkevicius M, Andersson DI, Sommer MOA. Collateral sensitivity constrains resistance evolution of the CTX-M-15 β-lactamase. Nature Communications. 2019;10(1):618.

30. Andrews JM, Howe RA. BSAC standardized disc susceptibility testing method (version 10). J. Antimicrob. Chemother. 2011;66(12):2726–57.

31. European Committee on Antimicrobial Susceptibility Testing. Breakpoint tables for interpretation of MICs and zone diameters. 2013;5.

32. García-Fernández S, Bala Y, Armstrong T, García-Castillo M, Burnham C-AD, Wallace MA, et al. Multicenter Evaluation of the New Etest Gradient Diffusion Method for Piperacillin-Tazobactam Susceptibility Testing of Enterobacterales, Pseudomonas aeruginosa, and Acinetobacter baumannii Complex. J. Clin. Microbiol. 2020;58(2):e01042–19.

33. European Committee on Antimicrobial Susceptibility Testing. Clinical Breakpoints - bacteria (v10.0). https://wwweucastorg/fileadmin/src/media/PDFs/EUCAST_files/Breakpoint_tables/v_100_Breakpoint_Tablespdf. 2020.

34. Bankevich A, Nurk S, Antipov D, Gurevich AA, Dvorkin M, Kulikov AS, et al. SPAdes: a new genome assembly algorithm and its applications to single-cell sequencing. J. Comput. Biol. 2012;19(5):455–77.

35. Seemann T. Prokka: rapid prokaryotic genome annotation. Bioinformatics. 2014;30(14):2068–9.

36. Hunt M, Mather AE, Sánchez-Busó L, Page AJ, Parkhill J, Keane JA, et al. ARIBA: rapid antimicrobial resistance genotyping directly from sequencing reads. Microb. Genom. 2017;3(10):e000131–e.

37. Inouye M, Dashnow H, Raven L-A, Schultz MB, Pope BJ, Tomita T, et al. SRST2: Rapid genomic surveillance for public health and hospital microbiology labs. Genom. Med. 2014;6(11):90.

38. Larsen MV, Cosentino S, Rasmussen S, Friis C, Hasman H, Marvig RL, et al. Multilocus sequence typing of total-genome-sequenced bacteria. J. Clin. Microbiol. 2012;50(4):1355–61.

39. Carattoli A, Zankari E, Garcia-Fernandez A, Voldby Larsen M, Lund O, Villa L, et al. In silico detection and typing of plasmids using PlasmidFinder and plasmid multilocus sequence typing. Antimicrob. Agents Chemother. 2014;58(7):3895–903.

40. Page AJ, Cummins CA, Hunt M, Wong VK, Reuter S, Holden MTG, et al. Roary: rapid large-scale prokaryote pan genome analysis. Bioinformatics. 2015;31(22):3691–3.

41. Page AJ, Taylor B, Delaney AJ, Soares J, Seemann T, Keane JA, et al. SNP-sites: rapid efficient extraction of SNPs from multi-FASTA alignments. Microb. Genom. 2016;2(4):e000056.

42. Nguyen LT, Schmidt HA, von Haeseler A, Minh BQ. IQ-TREE: a fast and effective stochastic algorithm for estimating maximum-likelihood phylogenies. Molecular biology and evolution. 2015;32(1):268–74.

43. Letunic I, Bork P. Interactive tree of life (iTOL) v3: an online tool for the display and annotation of phylogenetic and other trees. Nucleic Acids Res. 2016;44(W1):W242–5.

44. Croucher NJ, Page AJ, Connor TR, Delaney AJ, Keane JA, Bentley SD, et al. Rapid phylogenetic analysis of large samples of recombinant bacterial whole genome sequences using Gubbins. Nucleic Acids Res. 2015;43(3):e15.

45. Kallonen T, Brodrick HJ, Harris SR, Corander J, Brown NM, Martin V, et al. Systematic longitudinal survey of invasive Escherichia coli in England demonstrates a stable population structure only transiently disturbed by the emergence of ST131. Genome Res. 2017;27(8):1437–49.

46. Peter-Getzlaff S, Polsfuss S, Poledica M, Hombach M, Giger J, Böttger EC, et al. Detection of AmpC beta-lactamase in Escherichia coli: comparison of three phenotypic confirmation assays and genetic analysis. J. Clin. Microbiol. 2011;49(8):2924–32.

47. McNally A, Kallonen T, Connor C, Abudahab K, Aanensen DM, Horner C, et al. Diversification of Colonization Factors in a Multidrug-Resistant Escherichia coli Lineage Evolving under Negative Frequency-Dependent Selection. mBio. 2019;10(2):e00644–19.

48. Whitmer GR, Moorthy G, Arshad M. The pandemic Escherichia coli sequence type 131 strain is acquired even in the absence of antibiotic exposure. PLOS Pathogens. 2019;15(12):e1008162.

49. Cunha MPV, Saidenberg AB, Moreno AM, Ferreira AJP, Vieira MAM, Gomes TAT, et al. Pandemic extra-intestinal pathogenic Escherichia coli (ExPEC) clonal group O6-B2-ST73 as a cause of avian colibacillosis in Brazil. PLOS ONE. 2017;12(6):e0178970.

50. Rothe K, Wantia N, Spinner CD, Schneider J, Lahmer T, Waschulzik B, et al. Antimicrobial resistance of bacteraemia in the emergency department of a German university hospital (2013–2018): potential carbapenem-sparing empiric treatment options in light of the new EUCAST recommendations. BMC Infect. Dis. 2019;19(1):1091.

51. Baker TM, Rogers W, Chavda KD, Westblade LF, Jenkins SG, Nicolau DP, et al. Epidemiology of Bloodstream Infections Caused by Escherichia coli and Klebsiella pneumoniae That Are Piperacillin-Tazobactam-Nonsusceptible but Ceftriaxone-Susceptible. Open Forum Infect Dis. 2018;5(12):ofy300.

52. Stainton SM, Monogue ML, Nicolau DP. In Vitro-In Vivo Discordance with Humanized Piperacillin-Tazobactam Exposures against Piperacillin-Tazobactam-Resistant/Pan-β-Lactam-Susceptible Klebsiella pneumoniae Strains. Antimicrob Agents Chemother. 2017;61(7).

53. Carlisle L, Justo JA, Al-Hasan MN. Bloodstream Infection due to Piperacillin/Tazobactam Non-Susceptible, Cephalosporin-Susceptible Escherichia coli: A Missed Opportunity for De-Escalation of Therapy. Antibiotics. 2018;7(4).

54. Robson SE, Cockburn A, Sneddon J, Mohana A, Bennie M, Mullen AB, et al. Optimizing carbapenem use through a national quality improvement programme. J. Antimicrob. Chemother. 2018;73(8):2223–30.

55. Harrison EM, Ba X, Coll F, Blane B, Restif O, Carvell H, et al. Genomic identification of cryptic susceptibility to penicillins and β-lactamase inhibitors in methicillin-resistant Staphylococcus aureus. Nat. Microbiol. 2019;4(10):1680–91.

56. Williams CT, Edwards T, Adams ER, Feasey NA, Musicha P. ChloS-HRM, a novel assay to identify chloramphenicol-susceptible Escherichia coli and Klebsiella pneumoniae in Malawi. J. Antimicrob. Chemother. 2019.

57. Charalampous T, Kay GL, Richardson H, Aydin A, Baldan R, Jeanes C, et al. Nanopore metagenomics enables rapid clinical diagnosis of bacterial lower respiratory infection. Nat. Biotechnol. 2019;37(7):783–92.

58. Harmer CJ, Pong CH, Hall RM. Structures bounded by directly-oriented members of the IS26 family are pseudo-compound transposons. Plasmid. 2020;111:102530.

59. Sirot D, Chanal C, Bonnet R, De Champs C, Bret L. Inhibitor-resistant TEM-33 beta-lactamase in a Shigella sonnei isolate. Antimicrob. Agents Chemother. 2001;45(7):2179–80.

60. Che T, Bethel CR, Pusztai-Carey M, Bonomo RA, Carey PR. The different inhibition mechanisms of OXA-1 and OXA-24 beta-lactamases are determined by the stability of active site carboxylated lysine. J. Biol. Chem. 2014;289(9):6152–64.

61. Livermore DM, Day M, Cleary P, Hopkins KL, Toleman MA, Wareham DW, et al. OXA-1 β-lactamase and non-susceptibility to penicillin/β-lactamase inhibitor combinations among ESBL-producing Escherichia coli. J. Antimicrob. Chemother. 2018;74(2):326–33.

62. Nielsen KL, Hansen KH, Nielsen JB, Knudsen JD, Schønning K, Frimodt-Møller N, et al. Mutational change of CTX-M-15 to CTX-M-127 resulting in mecillinam resistant Escherichia coli during pivmecillinam treatment of a patient. MicrobiologyOpen. 2019;8(12):e941.

63. Edwards T, Williams C, Teethaisong Y, Sealey J, Sasaki S, Hobbs G, et al. A highly multiplexed melt-curve assay for detecting the most prevalent carbapenemase, ESBL and AmpC genes. Diag. Microbiol. Infect. Dis. 2020:115076.

64. Han MS, Park KS, Jeon JH, Lee JK, Lee JH, Choi EH, et al. SHV Hyperproduction as a Mechanism for Piperacillin-Tazobactam Resistance in Extended-Spectrum Cephalosporin-Susceptible Klebsiella pneumoniae. Microb. Drug. Resist. (Larchmont, NY). 2020;26(4):334–40.

65. Gilchrist CLM, Chooi Y-H. clinker & clustermap.js: automatic generation of gene cluster comparison figures. Bioinformatics. 2021;37(16):2473–5.

